# Exploring oxidative stress and endothelial dysfunction as a mechanism linking bisphenol S exposure to vascular disease in human umbilical vein endothelial cells and a mouse model of postnatal exposure

**DOI:** 10.1101/2022.08.15.504011

**Authors:** Sarah Easson, Radha Singh, Liam Connors, Taylor Scheidl, Larissa Baker, Anshul Jadli, Hai-Lei Zhu, Jennifer Thompson

**Author notes:** Authors contributed equally (Sarah Easson, Radha Singh). **Corresponding author:** Jennifer Thompson PhD, University of Calgary, Heritage Medical Research Building rm. 78, 3330 Hospital Dr. NW, Calgary, Alberta, Canada, T2N 4N1.

## Abstract

**Background:** Structural analogues used to replace bisphenol A (BPA) since the introduction of new regulatory restrictions are considered emerging environmental toxicants and remain understudied with respect to their biological actions and health effects. Studies reveal a link between BPA exposure and vascular disease in human populations, whereas the vascular effects of BPA substitutes remain largely unknown.

**Objectives:** To determine the effect of BPS, a commonly used BPA substitute, on redox balance, nitric oxide (NO) availability and microvascular NO-dependent dilation.

**Methods:** In human umbilical vein endothelial cells (HUVEC), production of reactive oxygen species (ROS) and NO after exposure to BPS was measured using fluorescent probes for DCFDA and DAF-FM diacetate, respectively. The contribution of endothelial NO synthase (eNOS) uncoupling to ROS generation was determined by measuring ROS in the presence or absence of an eNOS inhibitor (L-NAME) or eNOS co-factor, BH4, while the contribution of mitochondria-derived ROS was determined by treating cells with mitochondria-specific antioxidants prior to BPS exposure. Bioenergetic profiles were assessed using Seahorse extracellular flux analysis and mitochondria membrane polarization was measured with TMRE and JC-1 assays. In a mouse model of low dose BPS exposure, NO-mediated endothelial function was assessed in pressurized microvessels by inducing endothelium-dependent dilation in the presence or absence of L-NAME.

**Results:** BPS exposure (≥ 25 nM) reduced NO and increased ROS production in HUVEC, the latter corrected by treating cells with L-NAME or BH4. BPS exposure led to a loss of mitochondria membrane potential but had no impact on bioenergetic parameters except for a decrease in the spare respiratory capacity. Treatment of HUVEC with mitochondria-specific antioxidants abolished the effect of BPS on NO and ROS. NO-mediated vasodilation was impaired in male mice exposed to BPS.

**Discussion:** Exposure to BPS may promote cardiovascular disease by perturbing NO-mediated vascular homeostasis through the induction of oxidative stress.

## Introduction

The Lancet Commission on Pollution and Health estimated that pollution cost 9 million premature deaths per year, 62% due to cardiovascular disease (CVD).^1,2^ One of the most common chemical pollutants in our environment is Bisphenol A (BPA), a plasticizer used in the manufacture of polycarbonate plastics and epoxy resins. In recent years, global production of BPA has reached 5-6 billion pounds annually according to the CDC and is projected to grow over the next five years. Human exposure occurs primarily through direct consumption of BPA monomers leached from food and beverage containers but can also occur through inhalation and skin absorption and indirectly through contamination of soil, water, and air.^3,4^ Following consumption, BPA is absorbed in the gastrointestinal tract, rapidly metabolized in the liver, and excreted in the urine. Despite its short half-life, BPA is detected in over 90% of urine and blood samples as exposure is ubiquitous and continuous.^5,6^ The heavy environmental and health burden of industrial BPA production has made it a target of international efforts to safeguard planetary and public health.

In response to new regulatory restrictions on the use and import of BPA-containing infant products introduced in the late 2000s in Canada, US and Europe, the plastic industry began to substitute BPA with structural analogues including bisphenol S (BPS), bisphenol F (BPF) and bisphenol AF (BPAF). Whether these new synthetic bisphenols are safe alternatives to their predecessor remains unknown because the ‘innocent until proven guilty’ principal of chemical regulation liberates manufacturers from the burden of proving safety before marketing and widespread production. BPA substitutes are considered emerging environmental contaminants and are now detected in over 80% of urine samples.^7,8^ There remains scant data on the biological properties and health effects of BPA analogues, and thus investigation is needed to determine if these new substitutes offer benefits from a public health perspective.

A large body of evidence demonstrates wide-ranging health effects of BPA, including perturbations in sexual development, reproductive dysfunction, endocrine disorders, and cancer.^3,9,10^ Adverse health effects of BPA and its analogues have largely been attributed to their action as endocrine disrupting chemicals (EDC), synthetic chemicals that interfere with endocrine signaling. Few studies have focused on the cardiovascular effects of BPA or its substitutes; however, data in human populations reveal a potential relationship between BPA exposure and hypertension, coronary artery disease, and atherosclerosis.^11–14^ Endothelial dysfunction is a key early pathophysiological event in diseases of the heart and blood vessels. In the current study, we determined the impact of BPS on vascular redox balance, nitric oxide (NO) availability and NO-mediated endothelial function.

## Methods

### Cell Culture

Ea.hy926 human umbilical vein endothelial cells (HUVEC, ATCC) were cultured in DMEM containing 10% fetal bovine serum (FBS), 1% HT Supplement and 1% penicillin streptomycin, at 37°C and 5% CO_2_ and passaged 5-10 times. Once 75% confluency was reached, HUVEC were starved in 2% FBS 24 hrs prior to being exposed to various concentrations (2.5 nM to 25 μM) of BPS (Sigma) dissolved in DMSO.

### Animal model

Male and female C57BL6 mice (Jackson Laboratories) were housed in cages absent of plastic enrichment, fed a phytoestrogen-low diet (Teklab 2020), and maintained on a 12-hr light-dark cycle. At the time of weaning, littermates were randomly assigned to BPS (250 nM) or vehicle (DMSO), administered through the drinking water in glass bottles (Lab Products LLC). The concentration of 250 nM in the drinking water reflects a human equivalent estimated daily intake (EDI) of approximately 8 nmol BPS/kg body weight/day. This dose falls below the current tolerable daily intake (TDI) of 18 nmol BPA/kg body weight/day and 219 nmol BPA/kg body weight/day set by the European Food Safety Authority (EFSA) and the US. Food and Drug Administration (FDA), respectively.^15^ Currently, there are no TDI guidelines set for BPA analogues such as BPS. At 12 weeks of age, glucose tolerance was assessed in fasted mice by taking serial measurements of blood glucose from the tail with a glucometer after IP injection of glucose (2g/kg body weight). Mice were induced with 5% isoflurane with oxygen (1L/min) before euthanizing by decapitation. Serum was collected from trunk blood and the mesentery collected for dissection of microvessels. All animal procedures were approved by the University of Calgary Animal Care Committee (protocol #: AC19-0006) and conducted in accordance with the Canadian Council on Animal Care Ethics and the ARRIVE guidelines.

### Detection of ROS

HUVEC cultured in 96-well black plates were incubated in a 25 μM solution of 2’7’-dichlorodihydrofluorescein diacetate (DCFDA, Sigma, D6883) at 37°C for 30 min, and subsequently exposed to various concentrations of BPS or vehicle. The ROS generator, menadione (Sigma) was used as a positive control. After 30 min of exposure, ROS production was quantified by measuring the relative fluorescent intensity (RFI) of DCFDA (485/535 nm) on a SpectraMax M2 plate reader (Molecular Devices) and subsequently normalized to protein concentration. Oxidative stress was assessed in HUVEC exposed to BPS for 24 or 48 hrs by staining with 5 μM CellROX Deep Red™ (Invitrogen, C10422) for 30 min at 37°C. CellROX (640/665 nm) staining was quantified by plate reader or flow cytometry using an Attune® Acoustic Focusing Cytometer (Thermo Fisher Scientific) after cells were dissociated from the culture dish using 5 mM EDTA in Hanks balanced salt solution (HBSS), centrifuged at 500g and resuspended in HBSS. Images of HUVEC stained with CellROX and the Hoechst 33342 nuclear stain were captured on a Nikon Ti Eclipse Widefield Microscope. Mitochondria-derived ROS were identified with MitoSOX Red™ (Invitrogen, M36008) after 30 min of BPS exposure and quantified by fluorescence spectroscopy (510/580 nm) and flow cytometry. Three biological replicates were used for each assay.

### Production of NO and eNOS uncoupling

Production of NO was measured in HUVEC exposed to BPS for 30 or 90 min by loading with a DAF-FM Diacetate probe (Invitrogen, D23844) for 45 min at 37°C and quantifying on a SpectraMax M2 plate reader or flow cytometer. The endothelial NO synthase (eNOS) inhibitor, L-N^G^-Nitro arginine (L-NAME, Sigma), was used as a positive control. To assess the contribution of uncoupled eNOS to excess ROS production, cells were pre-incubated in L-NAME or tetrahydrobipterin (BH4, Sigma), a cofactor that stabilizes the eNOS dimer, prior to measurement of acute ROS production by the DCFDA fluorescent probe. The contribution of mitochondria-derived ROS to NO availability and vascular oxidative stress was determined by incubating cells with mitochondria-targeted antioxidants, MitoTempol (500 nM, Cayman Chemical) or SS31 (100 nM), for 1 hr prior to exposing cells to BPS. Three biological replicates were used for each assay.

### Antioxidant activity

Superoxide dismutase (SOD) activity was assessed via a commercially available kit (Cayman Chemical). In brief, after 48 hrs exposure of HUVEC to 2.5 uM BPS, cells were detached, centrifuged and the pellet was homogenized in cold HEPES digestion buffer (1 mM EGTA, 210 mM mannitol, 70mM sucrose). The digested samples were centrifuged, and the supernatant stored at −80ºC until assaying. Samples and standards were plated in triplicate. A radical detector and xanthine oxidase were added according to the manufacturer’s instructions. Following 30 min incubation at room temperature, absorbance was quantified using on a plate reader at 450nm and normalized to protein concentration.

### Gene expression analysis

After 48 hrs exposure to BPS or vehicle, total RNA was isolated from HUVEC using a RNeasy Mini Kit (Qiagen) according to the manufacturer’s instructions. The concentration of RNA was quantified with an N50 Nanophotometer (Implen Inc.) and RNA integrity evaluated with a TapeStation RNA assay (Agilent). Following DNase treatment, cDNA was generated using a High-Capacity cDNA Reverse Transcription kit (Applied Biosystems). cDNA was mixed with Powerup SYBR green master mix (Applied Biosystems) and primer pairs were run in triplicates on a QuantStudio 5 Real-time PCR instrument (Applied Biosystems). Primers were designed using the NCBI/Primer Blast tool (Table 1) and tested for specificity using melting curve analysis. Target mRNA expression was normalized to ribosomal protein lateral stalk subunit (RPLP0) and fold change calculated using the 2^−ΔΔCt^ method. Expression data were further verified using beta-actin as a second internal control for normalization.

**Table 1:**
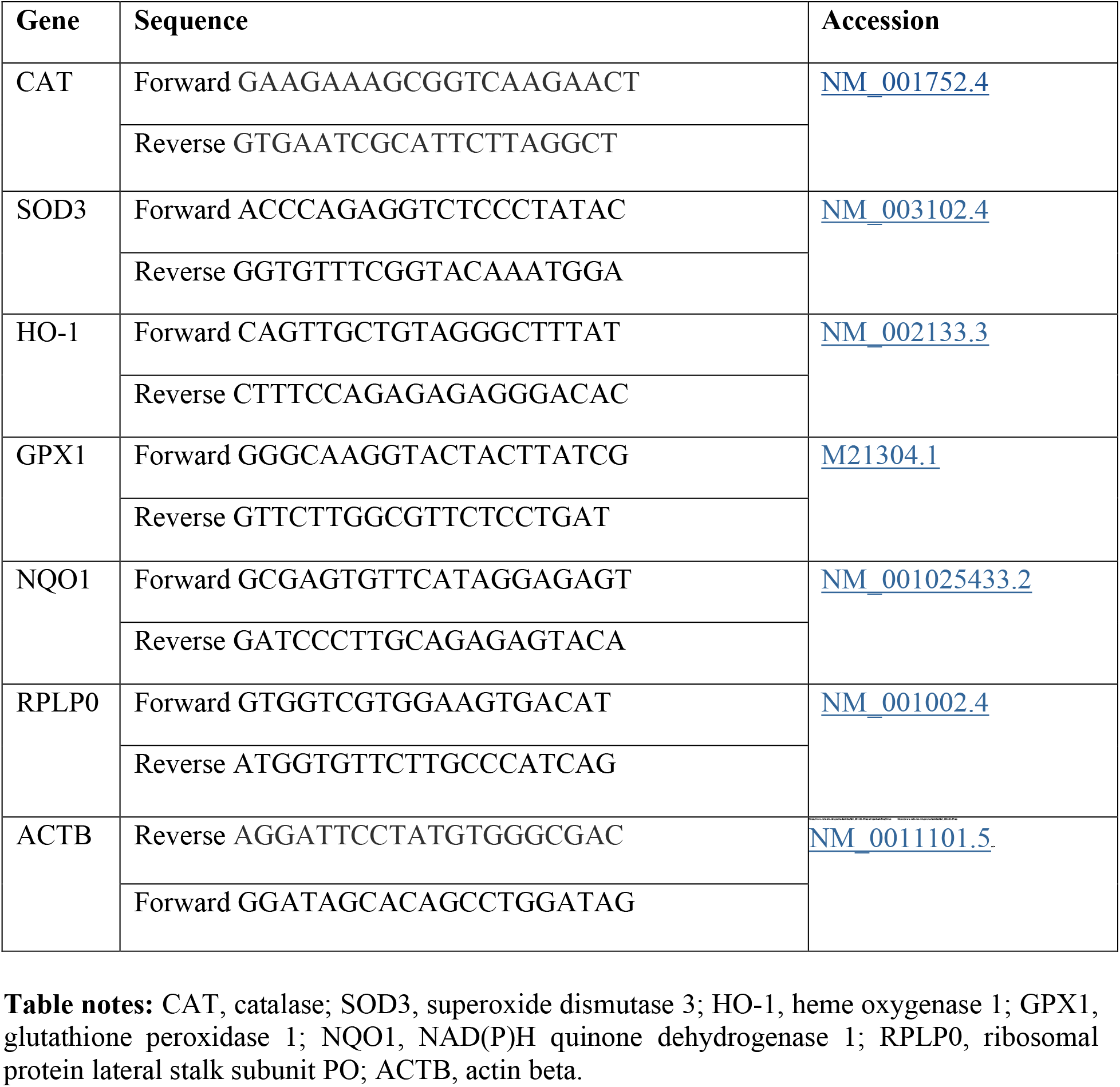
Primersrep

### Western blot

Whole cell lysates were homogenized in RIPA buffer (Invitrogen) and protein quantified with a BCA protein assay (Thermo Scientific). Protein (40 µg) was loaded into pre-casted NuPAGE 4-12% Bis-Tris gels (Invitrogen) and separated by electrophoresis. After transferring separated protein onto PVDF membranes (GE Healthcare Life Sciences), membranes were blocked in 5% (w/v) non-fat dry milk (Biorad) diluted in TBST for 1 hour. Membranes were probed overnight in primary antibodies [phospho-eNOS (Cell Signaling, rabbit 9571); eNOS (Cell Signaling, rabbit 32027); phospho-Akt (Cell Signaling, rabbit 4060); Akt (Cell Signaling, rabbit 4691); GAPDH (Cell Signaling, rabbit 2118) that were pre-incubated in the blocking buffer for 1 hour. Following washing of the membrane and incubation in secondary anti-rabbit antibody, immunoreactive proteins were imaged on an iBright 1500 Imaging System (Molecular Probes) using enhanced chemiluminescence (ECL).

### Mitochondria membrane potential

HUVEC were exposed to BPS for 48 hrs and assessed for mitochondria function. Mitochondria membrane polarization was measured by loading cells with 2 µM JC-1 (5, 5’, 6, 6’-tetrachloro-1, 1’, 3, 3’-tetraethylbenzimidazolylcarbocyanine iodide, Invitrogen, 15003) at 37°C for 15 min. After washing, fluorescence was measured at 514/529 (green, monomer) and 514/590 (red, aggregate) using the SpectraMax M2 plate reader. The mitochondria membrane potential was also assessed in cells stained with 50 nM tetramethyl rhodamine ethyl ester (TMRE, ThermoFisher, T669) and subsequently collected in HBSS containing 5 mM EDTA, washed and resuspended in media. The TMRE fluorescent signal was quantified on a plate reader and flow cytometer. The mitochondria uncoupler, FCCP, was used as a positive control. Three biological replicates were used for each assay.

### Mitochondria bioenergetics

Extracellular flux analysis was performed with HUVEC exposed to 2.5 µM BPS or vehicle using a Seahorse XFp Analyzer (Agilent Technologies). Cells were plated at 5000 cells/well on Seahorse XFp miniplates. After 48 hrs of exposure, cells were switched to Seahorse XF assay media (non-buffered DMEM containing 2 mM L-glutamine, 1mM sodium pyruvate and 10 mM glucose) and incubated for 45 min. The oxygen consumption rate (OCR), extracellular acidification rate (ECAR) and other bioenergetic parameters were measured using the Agilent Seahorse XFp Mito stress test kit (Agilent, 103010-100). Baseline and stressed OCR and ECAR were determined prior and post-injection of stressor compounds, oligomycin (1.5 µM), FCCP (0.5 µM) and rotenone/antimycin A (0.5 µM), respectively. The data were normalized to protein content from individual wells. Data were analyzed using Wave software version 2.6 (Agilent Technologies).

### Isolation of microvessels and arterial pressure myography

Fourth order mesenteric arteries were carefully excised from the mesentery under a dissecting microscope and cleared of perivascular fat in ice-cold Krebs solution (120 mM NaCl, 25mM NaHCO_3_, 4.8 mM KCl, 11 mM glucose, 0.27 mM EDTA, 1.2 mM NaH_2_PO_4_, 1.2 mM of MgSO_4_, and 2.5 mM CaCl_2_; pH 7.4). The vessels were mounted and secured with sutures onto glass cannulas diametrically opposed in a pressure myograph chamber (Living Systems), which was placed on the stage of an inverted microscope (Nikon) equipped with live video recording. The pressure myograph chamber was aerated with 5% CO_2_/95% O_2_ air gas mixture and cannulated vessels maintained at 37°C at a constant intraluminal pressure of 60 mmHg under no flow conditions with a servo-controlled pressure device (Living Systems). Vessel diameter was recorded using IonOptix software. After equilibration for 30 min, arterial viability was assessed by stimulation with a dose of saturated KCL solution. Contractile responses were assessed by adding cumulative concentrations (10^−10^ M to 10^−4^ M) of the α-adrenergic agonist, phenylephrine (Phe). After pre-contraction with 10^−6^ M Phe, NO-dependent dilation was determined by generating concentration response curves to methacholine (Mch, 10^−11^ M to 10^−5^ M). To determine the contribution of NO to Mch-induced dilation, a subset of vessels was incubated with L-NAME for 30 min prior to generating Mch concentration-response curves. Endothelium-independent dilation was assessed by measuring dilation in response to 10^−4^ M sodium nitroprusside in pre-contracted vessels. After concentration response curves were generated, the Krebs solution was replaced with Ca^2+^-free buffer and equilibrated for 45 min before recording passive diameter at 60 mmHg. Contractile and dilatory responses were normalized to passive diameter and expressed as a percentage of maximum KCL-induced contraction or pre-contraction, respectively.

### Statistical analyses

Statistical analyses were carried out using GraphPad Prism version 9.3.1. Two-tailed student’s t-test or One-way ANOVA with Dunnett’s or Tukey’s multiple comparison test were used to evaluate differences between groups for *in vitro* experiments. Two-way ANOVA was performed to evaluate differences in concentration response curves. For calculation of the EC_50_, concentrations were converted to log form, data were normalized and the EC_50_ was compared by a nonlinear regression using the least squares method. Differences were considered statistically significant if p < 0.05 and data are presented as mean ± SEM.

## Results

### ROS production in HUVEC exposed to BPS

Production of ROS was increased in HUVEC acutely exposed to 25 nM to 25 µM BPS for 30 min (Figure 1A). After 24 and 48 hrs of BPS exposure, an increase in CellROX staining was observed at BPS concentrations as low as 2.5 nM (Figure 1B-E). HUVEC exposed to BPS exhibited decreased mRNA expression of key genes involved in antioxidant defences against oxidative stress (Figure 1F), including superoxide dismutase 3 (SOD3), heme oxygenase (HO-1), glutathione peroxidase 1 (GPX1), and NADPH quinone 1 dehydrogenase (NQO1). The activity of SOD was reduced in HUVEC exposed to 2.5 µM for 48 hrs (Figure 1G).

**Figure 1:**
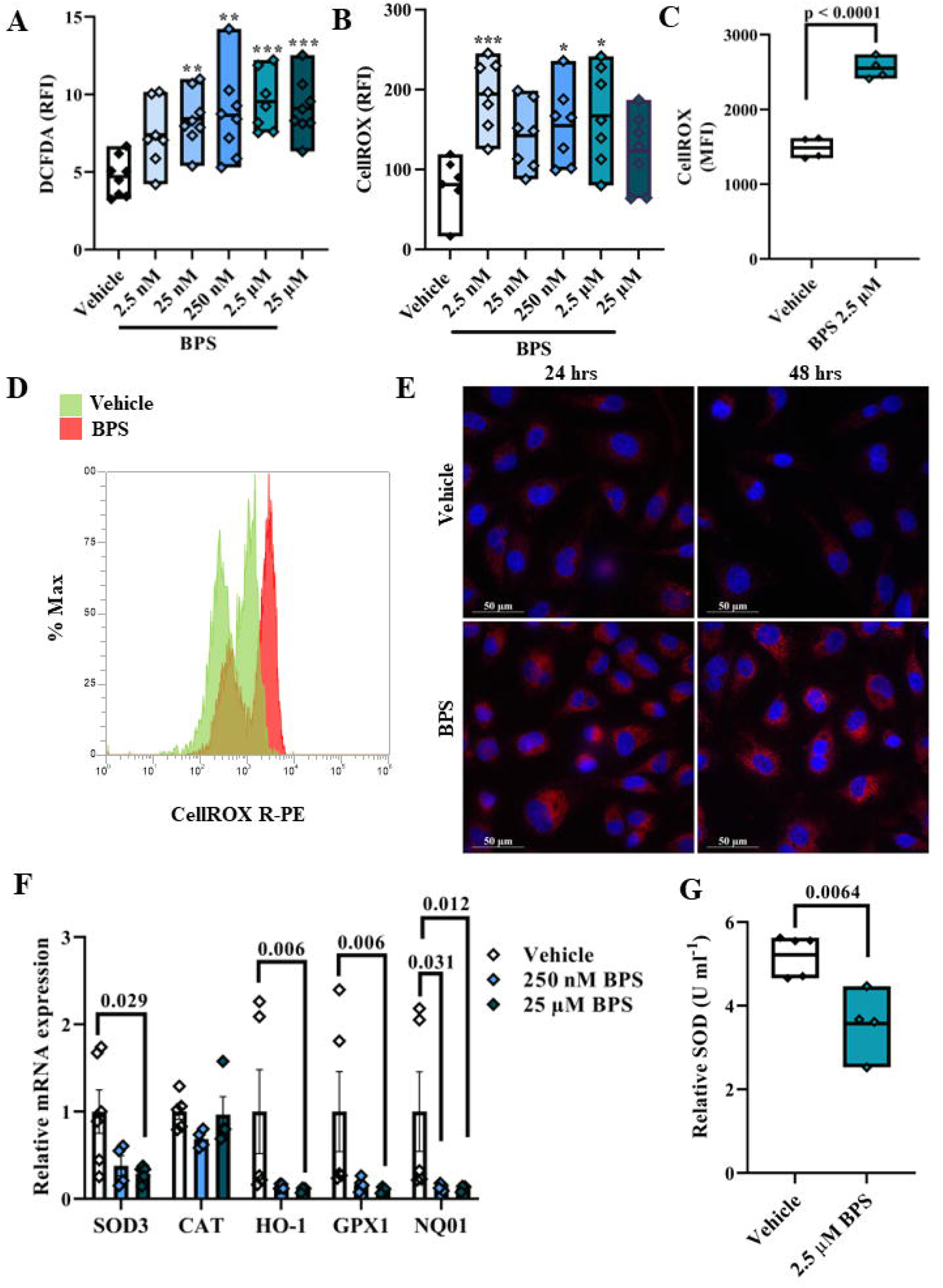
Redox balance in HUVEC exposed to BPS. Production of ROS was measured by quantifying relative fluorescent intensity (RFI) by fluorescence spectroscopy in HUVEC exposed to BPS for 30 min and loaded with a DCFDA probe (A). After 24 or 48 hrs of BPS exposure, CellROX Deep Red stain, an indicator of oxidative stress, was quantified by fluorescence spectroscopy (B). CellROX staining was also quantified using flow cytometry (C & D) and expressed as mean fluorescent intensity (MFI). Images of CellROX-stained HUVEC were captured on an epifluorescence-equipped microscope (E). mRNA expression analysis of genes involved in antioxidant defences was performed by qPCR (F) and the activity of superoxide dismutase (SOD) was determined using a SOD activity assay kit (G). Differences were compared using one-way ANOVA with a Dunnett’s multiple comparison test. * p < 0.05; ** p < 0.01; *** p < 0.0001 BPS vs. vehicle.

### NO availability and the contribution of eNOS uncoupling in BPS-exposed HUVEC

When the production of the free radical, superoxide (O_2_^−^) exceeds antioxidant capacity, O_2_^−^ reacts with NO to reduce its bioavailability. NO production in HUVEC was reduced when exposed to BPS for 30 min (Figure 2A) or 90 min (Figure 2B). The phosphorylation of eNOS at serine 1177 was reduced in HUVEC exposed to 2.5 µM BPS for 48 hrs, while BPS had no impact on phosphorylation of Akt at serine 473 or total eNOS expression (Figure C & D). Under conditions of oxidative stress, eNOS becomes uncoupled from L-arginine oxidation and produces O_2_^−^ rather than NO, contributing to a vicious cycle of excess ROS production and NO depletion. In HUVEC exposed to BPS for 48 hrs, inhibition of eNOS with L-NAME abolished the observed differences in DCFDA staining between BPS and vehicle (Figure 2E & G). BH4 is a critical cofactor that stabilizes the active eNOS dimer and its oxidation by ROS contributes to eNOS uncoupling. Supplementation with BH4 decreased DCFDA staining in BPS-exposed HUVEC but had no impact on control cells (Figure 2F & G), providing further evidence for eNOS uncoupling. Thus, heightened ROS production and uncoupled eNOS in BPS-exposed endothelial cells lead to reduced bioavailability of NO.

**Figure 2:**
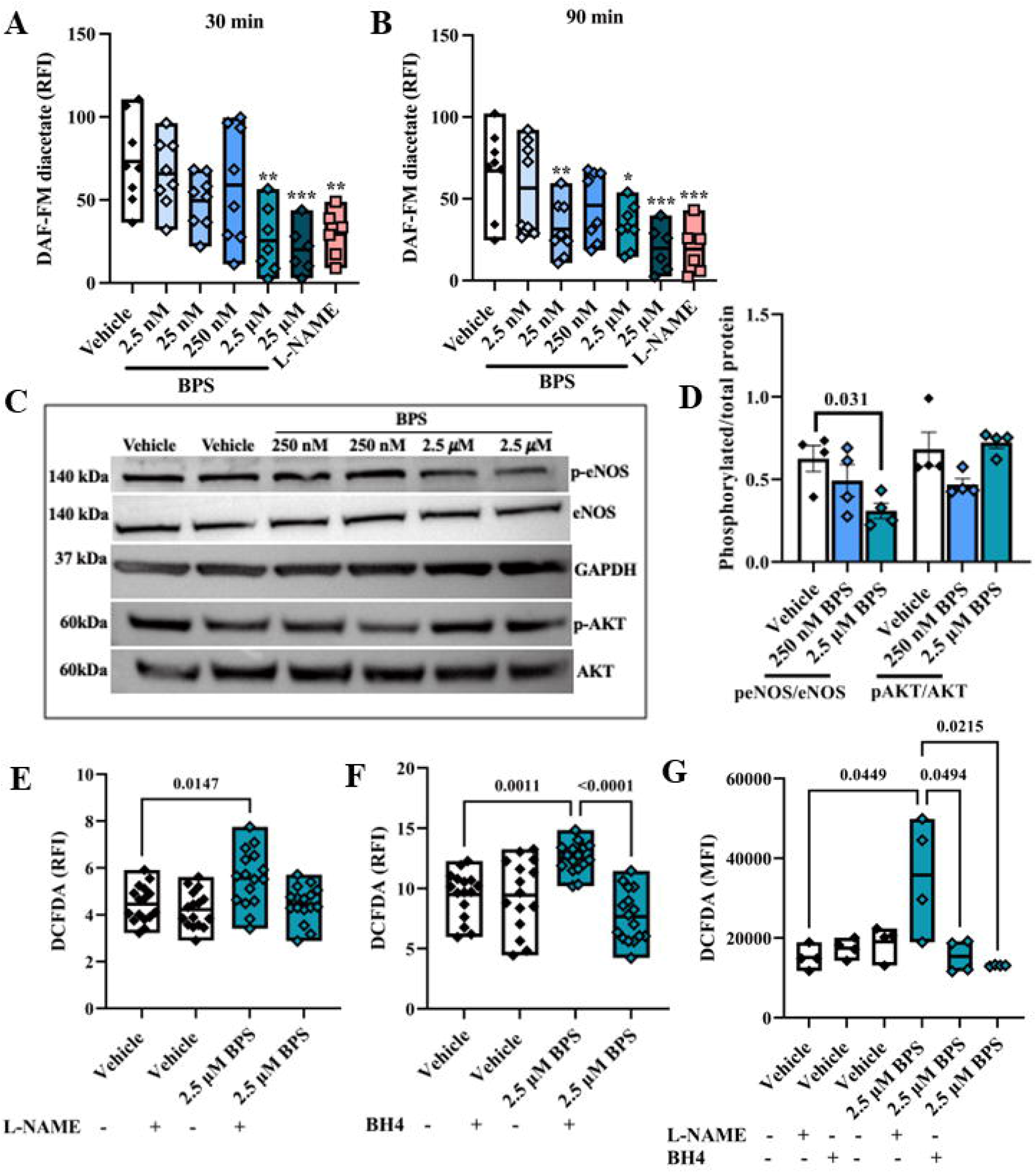
NO production and eNOS uncoupling in HUVEC exposed to BPS. Relative fluorescence intensity (RFI) of DAF-FM diacetate, a fluorescent NO probe, was measured in HUVEC exposed to BPS for 30 (A) or 90 min (B). Representative blots showing expression of phosphorylated and total eNOS or AKT in HUVEC exposed to BPS for 48 hrs (C). The ratio of phosphorylated/total protein was calculated by densitometry (D). The contribution of eNOS to ROS production was determined by inhibiting eNOS with L-NAME or supplementing with BH4 prior to quantification of DCFDA. DCFDA fluorescence was quantified by spectroscopy (E & F) or flow cytometry (G). Comparisons were made by one-way ANOVA with a Dunnett’s post hoc test (A-D, BPS vs. vehicle) or Tukey’s post hoc test (E-G). * p < 0.05; ** p < 0.01; *** p < 0.001.

### The contribution of mitochondria function to NO availability in BPS-exposed HUVEC

Dysfunctional mitochondria are a major source of excess ROS and contribute to the oxidative stress that drives endothelial dysfunction. A loss of membrane potential in BPS-exposed HUVEC was apparent by lower formation of J-aggregates when JC-1 enters the mitochondria, relative to cationic JC-1 monomers, (Figure 3A & B). Likewise, lower accumulation of the cationic dye, TMRE (Figure C-E) indicates mitochondria membrane depolarization. Mitochondria-derived ROS were increased in HUVEC exposed to 25 nM to 25 µM of BPS (Figure 3F-H). Figure 4 shows bioenergetic parameters measured by extracellular flux analysis in HUVEC exposed to BPS for 48 hrs. There were no differences in basal respiration or maximal respiration, but spare respiratory capacity was reduced in BPS-exposed HUVEC (Figure 4E), suggesting a compromised ability of mitochondria to respond to stress. Treating cells with mitochondria-targeted antioxidants prior to BPS exposure rescued NO production and prevented oxidative stress (Figure 5), suggesting that mitochondria-derived ROS contribute importantly to perturbed redox balance and NO signaling in BPS-exposed endothelial cells.

**Figure 3:**
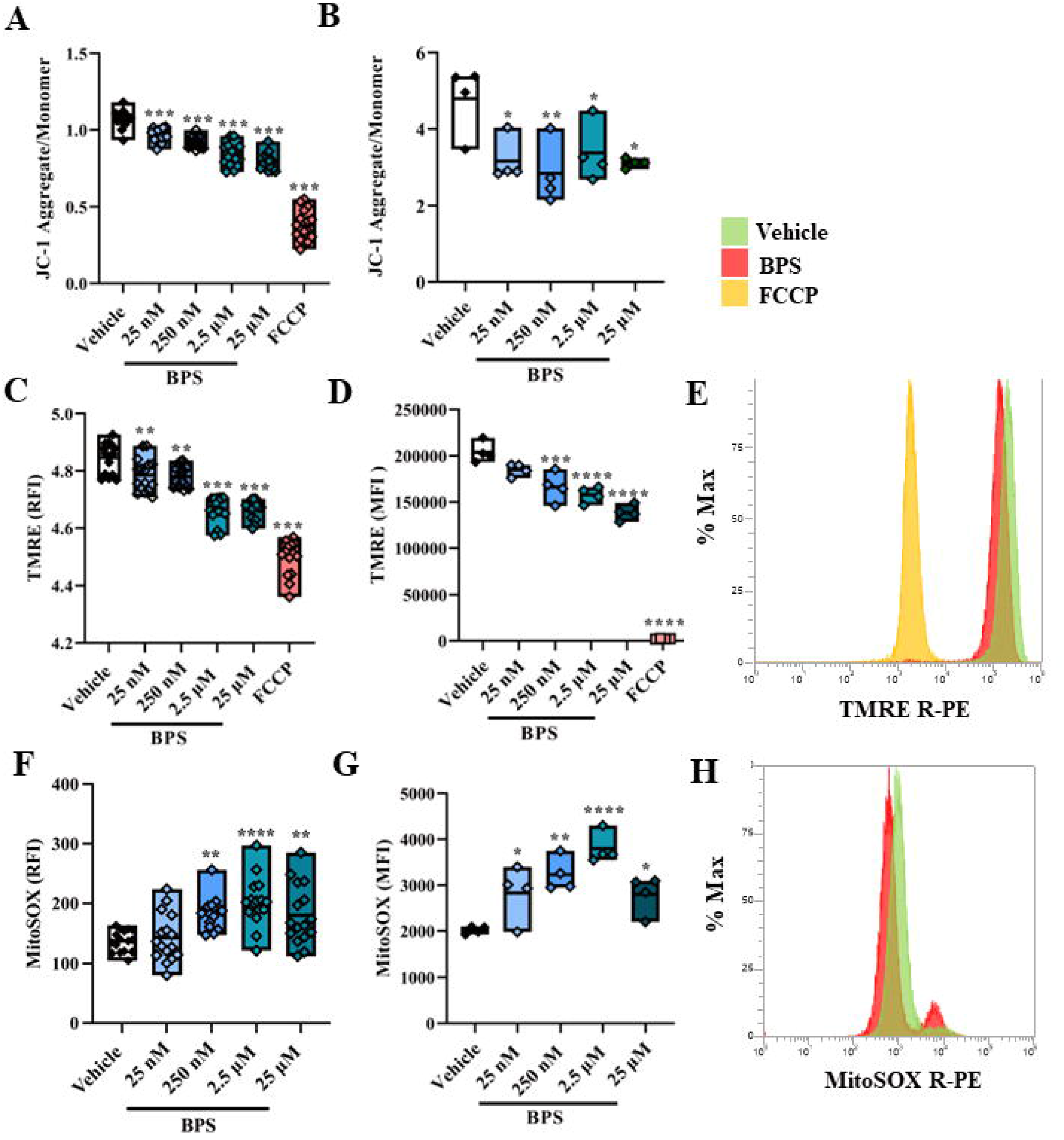
Mitochondria membrane polarization and mitochondria-derived ROS in HUVEC exposed to BPS. Mitochondria polarization was determined by calculating the ratio of red fluorescence (JC-1 aggregates) to green fluorescence (JC-1 monomers) using either fluorescent spectroscopy (A) or flow cytometry (B). Mitochondria membrane potential was also determined by quantifying fluorescence of the cationic dye, TMRE, by spectroscopy (C) or flow cytometry (E & F). Mitochondria-derived ROS was measured in HUVEC exposed to BPS for 30 min by measuring the fluorescence of MitoSOX Red by plate reader (F) or flow cytometric methods (G & H). Differences were compared by one-way ANOVA with a Dunnett’s multiple comparison test. * p < 0.05; ** p < 0.01; *** p > 0.001 BPS vs. vehicle.

**Figure 4:**
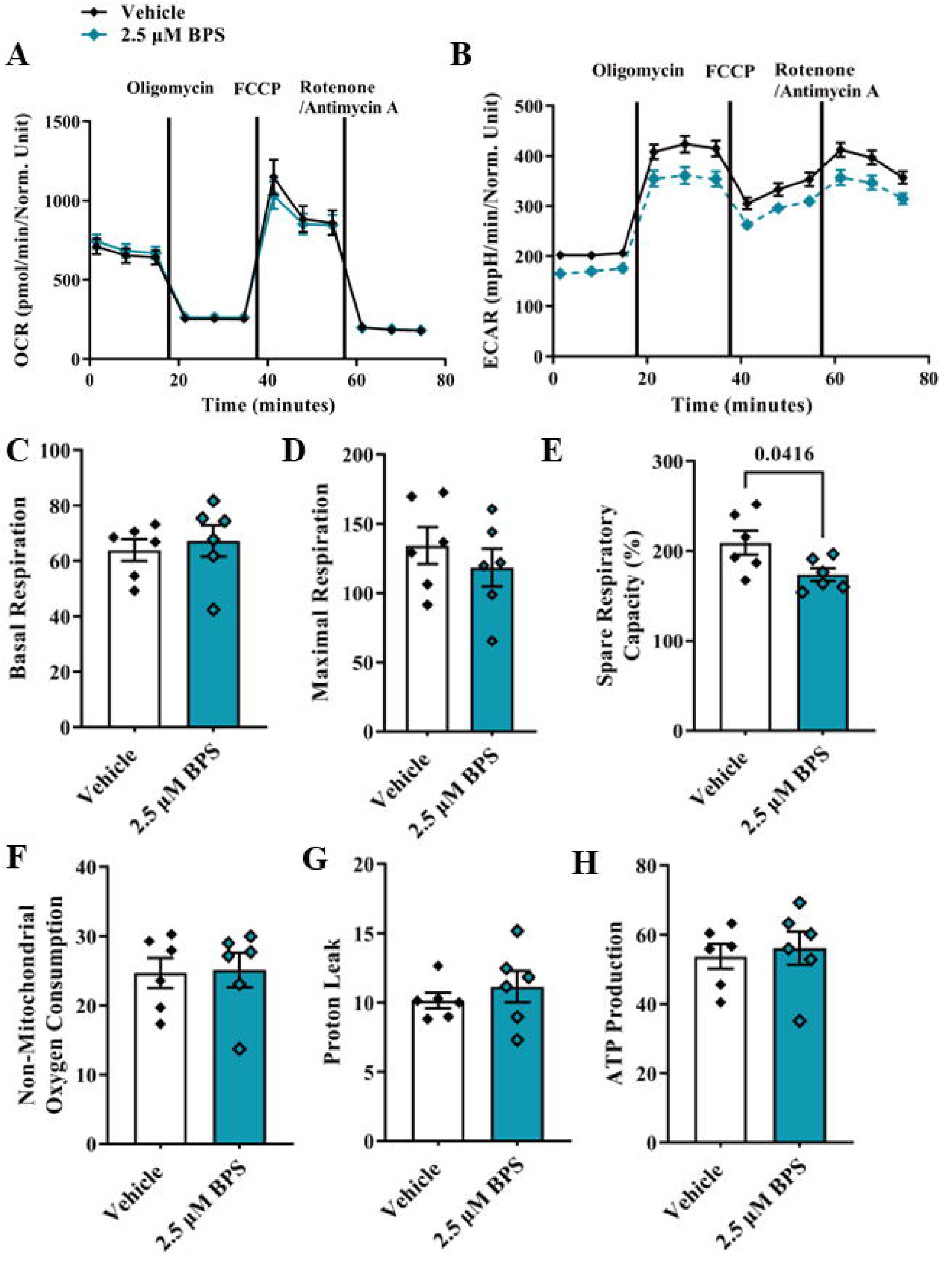
Bioenergetic profile of HUVEC exposed to BPS. Extracellular flux analysis was performed with a Seahorse Xfp Analyzer using a Mito stress test kit. After 48 hrs of exposure to 2.5 µM BPS, the oxygen consumption rate (OCR) (A) and extracellular acidification rate (ECAR) (B) were measured under basal conditions and after injection of oligomycin, FCCP and rotenone/antimycin A. Basal respiration (C), maximal respiration (D), spare respiratory capacity (E), non-mitochondrial oxygen consumption (F), proton leak (G) and ATP production (H) were calculated. Differences were compared with a student’s t-test.

**Figure 5:**
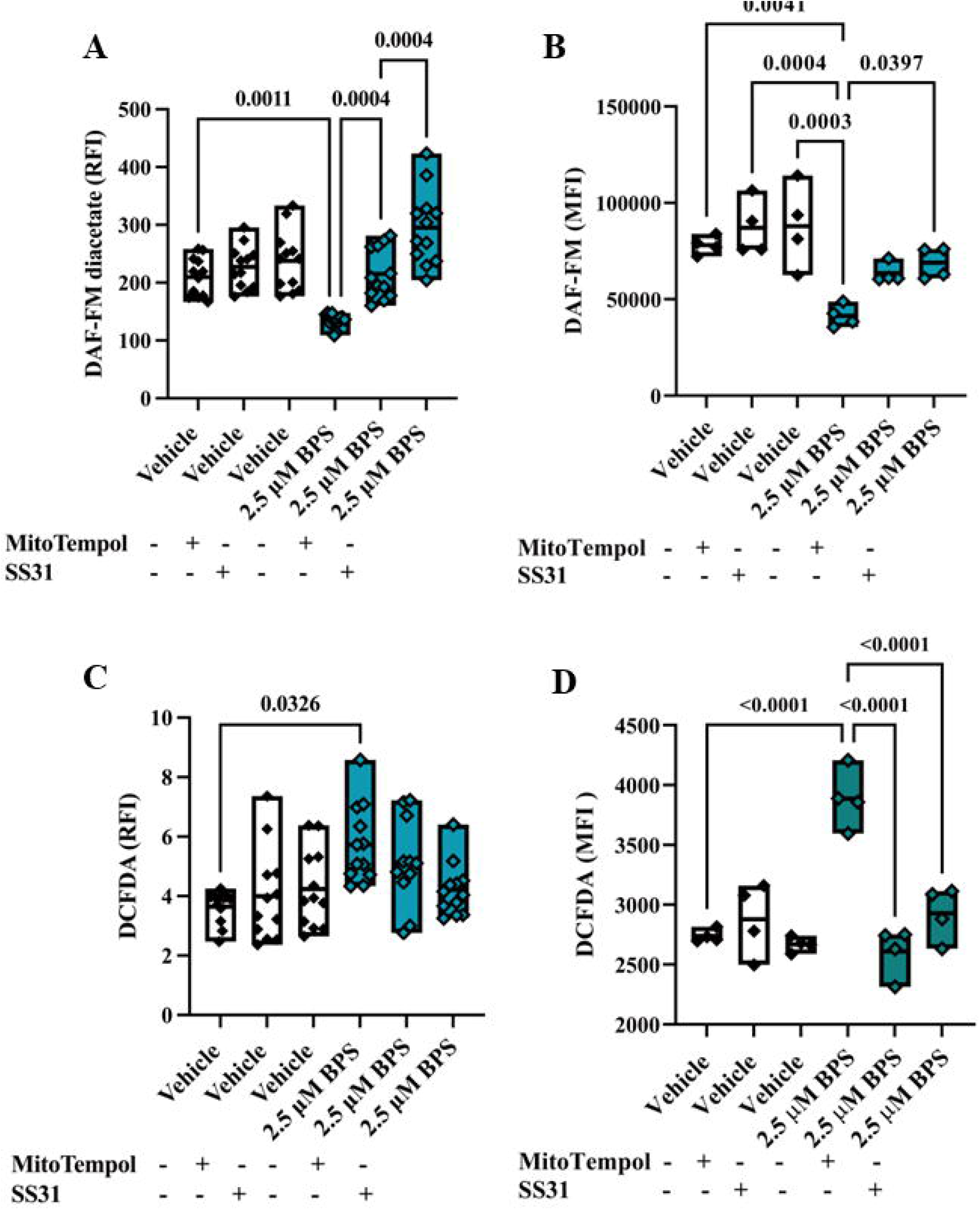
The contribution of mitochondria to oxidative stress and NO availability after BPS exposure. HUVEC were treated with the mitochondria-targeted antioxidants, MitoTempol or SS31, for 1 hr prior to BPS exposure. NO production was measured by loading cells with a DAF-FM diacetate probe (A & B) and ROS production was measured using a DCFDA probe (C & D). Fluorescence was quantified by spectroscopy and flow cytometry. Differences were compared using one-way ANOVA with Tukey’s post hoc test.

### Endothelium-dependent dilation in intact microvessels isolated from BPS-exposed mice

To determine if NO-dependent vasodilation is impaired in a mouse model of BPS exposure, male and female C57BL6 mice were exposed to the human equivalent of 8 nM BPS/kg body weight/day from 3 to 12 weeks of age. Endothelium-dependent dilation was assessed *ex vivo* in 4^th^ order mesenteric arteries by measuring diameter after adding cumulative doses of Mch after pre-contraction with Phe, while contractile responses were subsequently evaluated by adding cumulative doses of Phe (Supplementary Figure 1). BPS exposure had no impact on arterial contractile or dilatory responses in females (Figure 6A & B). Similar to females, there were no differences in contractile responses to the α1-adrenergic agonist, Phe, between control and BPS-exposed males (Figure 6C). However, microvessels isolated from BPS-exposed males exhibited reduced endothelium-dependent vasodilatory responses to the muscarinic agonist, Mch (Figure 6D & E). Endothelium-independent vasodilatory responses to the smooth muscle relaxant, SNP, were similar between groups. Pre-incubation with L-NAME blunted Mch responses in both vehicle and BPS-exposed males (Figure 6D & E). The NO-sensitive component of endothelium-dependent dilation was 63% in control males vs. 27% in BPS-exposed males, while the non-NO component of vasodilation was 37% in control and 73% in exposed males (Figure 6F). The EC_50_ values of Mch and Phe concentration response curves, as well as passive diameter and maximum responses to KCL are shown in Table 2. The passive diameter of mesenteric arteries measured in Ca^2+^-free conditions was decreased in BPS-exposed female mice, but not in BPS-exposed male mice. No differences in glucose tolerance were observed in either male or female mice exposed to. BPS (Supplementary Figure 2).

**Table 2:** EC50

|  |  | Sex | Vehicle | BPS |
| --- | --- | --- | --- | --- |
| Methacholine (Mch) | EC<sub>50</sub> (M)<br>(95% CI) | Male | 6.99x10<sup>-9</sup><br>(3.24x10<sup>-9</sup>, 1.47x10<sup>-8</sup>) | 9.49x10<sup>-9</sup><br>(2.93x10<sup>-9</sup>, 2.88x10<sup>-8</sup>) |
| Methacholine (Mch) | EC<sub>50</sub> (M)<br>(95% CI) | Female | 5.20x10<sup>-8</sup><br>(1.79x10<sup>-8</sup>, 1.44x10<sup>-7</sup>) | 1.97x10<sup>-8</sup><br>(4.50x10<sup>-9</sup>, 8.19x10<sup>-8</sup>) |
| Phenylephrine (Phe) | EC<sub>50</sub> (M)<br>(95% CI) | Male | 5.10x10<sup>-7</sup><br>(1.58x10<sup>-7</sup>, 2.01x10<sup>-6</sup>) | 4.89x10<sup>-7</sup><br>(1.78x10<sup>-7</sup>, 2.87x10<sup>-6</sup>) |
| Phenylephrine (Phe) | EC<sub>50</sub> (M)<br>(95% CI) | Female | 3.57x10<sup>-7</sup><br>(1.60x10<sup>-7</sup>, 7.89x10<sup>-7</sup>) | 4.71x10<sup>-7</sup><br>(1.21x10<sup>-7</sup>, 1.79x10<sup>-6</sup>) |
| KCl | Contraction<br>(% ± SEM) | Male | 40.9 ± 0.90 | 42.7 ± 3.51 |
| KCl | Contraction<br>(% ± SEM) | Female | 43.0 ± 3.01 | 39.3 ± 3.69 |
| Passive (Ca<sup>2+</sup>) diameter | Diameter<br>(µm± SEM) | Male | 216.7 ± 12.02 | 239.1 ± 6.99 |
| Passive (Ca<sup>2+</sup>) diameter | Diameter<br>(µm± SEM) | Female | 226.8 ± 9.69 | 193.5 ± 10.89<br>* p = 0.036 |

**Table notes:** Half maximal effective concentration (EC<sub>50</sub>) of methacholine (Mch) or phenylephrine (Phe) concentration-response curves generated in mesenteric arteries were calculated by transforming concentrations to log form, normalizing the data and fitting to nonlinear least squares fit curve. Maximal responses to a high dose of potassium chloride (KCl) were calculated relative to baseline. Passive internal diameter was recorded in pressurized mesenteric arteries under Ca<sup>2+</sup>-free conditions and compared by student's t-test. n = 6-15/group

**Figure 6:**
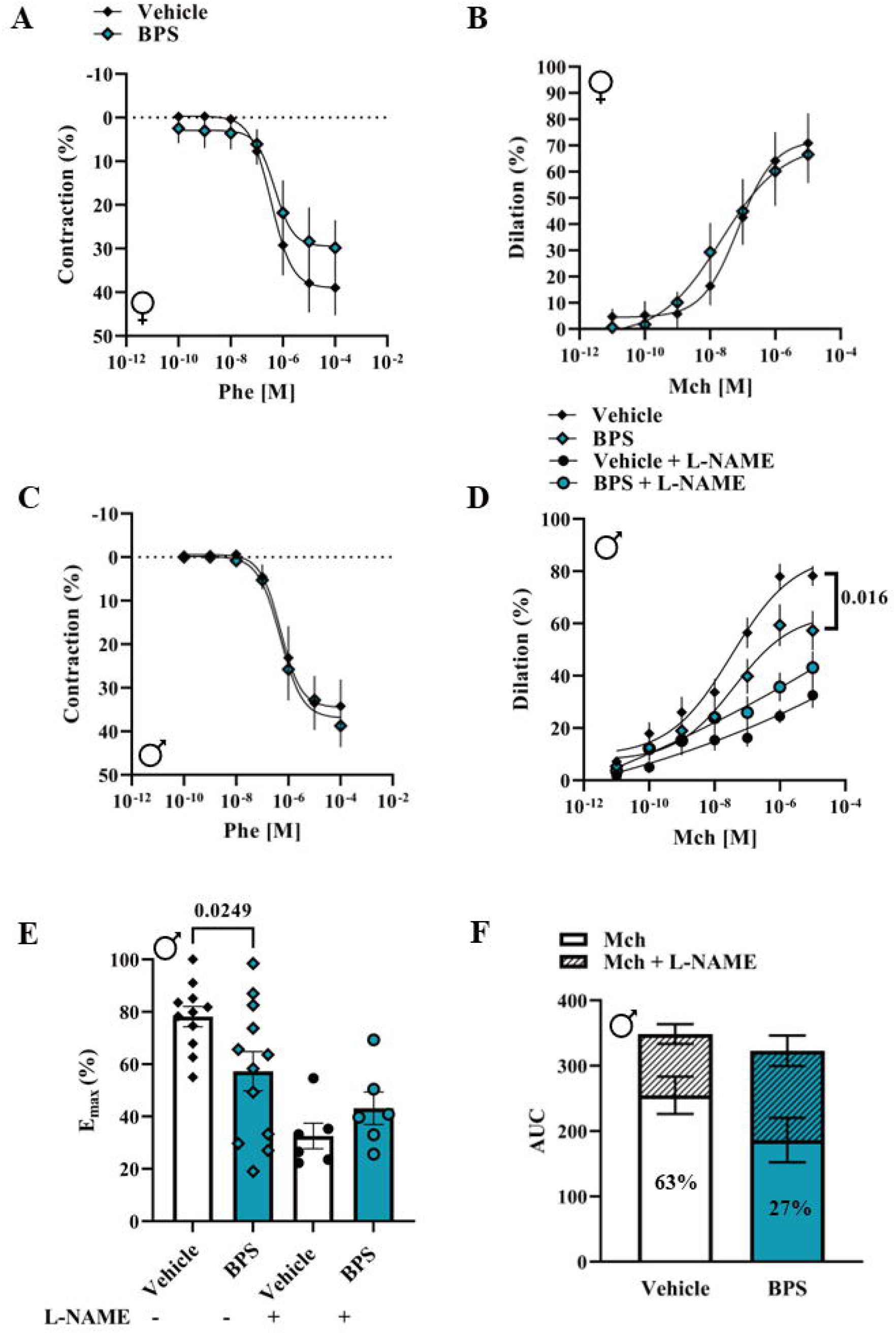
Endothelium-dependent vasodilation in microvessels isolated from male and female mice exposed to BPS. Concentration-dependent contractile responses to phenylephrine (Phe) were measured in pressurized mesenteric arteries from female (A) and male (C) C57BL/J6 mice exposed to vehicle or 250 nM BPS via drinking water for ~ 9 weeks. Responses to cumulative concentrations of the endothelium-dependent vasodilator, methacholine (Mch), were calculated in pressurized Phe pre-contracted arteries isolated from female (B) or male (D) mice. In a subset of male vessels, concentration-dependent dilatory responses to Mch were assessed in the presence of the eNOS inhibitor, L-NAME (D). The % change in diameter was normalized to passive diameter under Ca^2+^-free conditions, concentrations were log-transformed, fitted to a non-linear least squares curve, and compared by two-way ANOVA. The p value denotes significance in the interaction between concentration of Mch and treatment group. E_max_ of Mch responses in the presence or absence of L-NAME were compared by two-way ANOVA (Figure E). Area under the curve (AUC) was calculated and the AUC of dilatory responses in the presence of L-NAME subtracted from the Mch curve without L-NAME to determine the NO-sensitive component of vasodilation in vehicle (63%) and BPS-treated males (27%). n = 6-15 mice/group.

## Discussion

Structural analogues of BPA are increasingly used by manufacturers and marketed as safer substitutes, yet their biological actions and health effects remain largely unknown. Findings from the current study show that exposure to BPS, a common substitute for BPA, leads to sex-dependent impairments in microvascular function and reveal reduced NO availability due to oxidative stress as a probable underlying mechanism. Endothelium-dependent vasodilatory responses were impaired in intact microvessels isolated from male mice exposed to BPS, while BPS exposure had no impact on microvascular endothelial function in female mice. Pre-incubating vessels with the eNOS inhibitor, L-NAME, revealed a shift to higher contributions of NO-independent signaling in BPS-exposed males, suggesting a compensatory recruitment of other vasodilators such as prostaglandin. Production of NO by the vascular endothelium is not only critical in maintaining vascular tone and blood pressure, but plays a key role in promoting vascular homeostasis as NO exhibits anti-inflammatory, anti-thrombotic and anti-atherogenic properties.^16,17^ Blunted NO production is a hallmark of endothelial dysfunction, a common pathophysiological mechanism in the development of hypertension, atherosclerosis and other diseases of the heart and vasculature.^17^ Epidemiological studies using data from the National Health and Nutrition Examination Survey (NHANES) show that urinary BPA levels predict the risk for hypertension, coronary artery disease and other CVD.^13,18^ Far fewer studies have focused on the new structural analogues of BPA. Recently, cross-sectional studies examining populations in China and the USA reported positive associations between BPA and BPS exposure with hypertension or CVD, but no effect of BPF, another common substitute for BPA.^19,20^ Our study reveals endothelial dysfunction as a key mechanism linking BPS to the development of CVD. To our knowledge, no other study has examined the effect of *in vivo* BPS exposure on endothelial function in intact vessels; however, a study by Saura *et al*. showed that exposure of mice to 400 nM BPA through the drinking water increased systolic and diastolic blood pressure and impaired endothelium-dependent relaxation in carotid arteries.^21^ The sex of mice used in the above-mentioned study is not noted; therefore, it is unknown if sex-dependent vascular effects are also observed with BPA exposure. The current study shows that males are particularly susceptible to BPS-induced endothelial dysfunction, highlighting the importance of considering sex differences in the health effects of EDC.

The dose of BPS used for *in vivo* exposure to test the impact on endothelial function in intact microvessels was 250 nM in the drinking water, which is approximately a human equivalent of 1.9 µg/kg body weight/day or 8 nM/kg body weight/day. Although there are no TDI established for BPA analogues, the TDI for BPA currently set by the EFSA is 4 µg/kg body weight/day or 18 nM/kg body weight per day.^22^ Recently published results from the Canadian Total Diet Study (TDS) reveal that estimated ingestion of BPS from contaminated foodstuffs, a major source of human exposure, ranged between 5.74 to 56.9 ng/kg body weight/day in the provinces of China, and exceeded that of BPA intake.^23^ In a cohort of pregnant women recruited for the APrON study in Alberta, Canada, daily 24 hr intake of BPS reached up to 14 nM/kg body weight, approaching the TDI of BPA.^24^ It is well established that bisphenol exposure is higher in young children; ingestion rates have been reported to be ~ 4-fold higher in toddlers compared to adults.^25^ Therefore, the dose used in the present study is below the TDI and is environmentally relevant with respect to human exposure. Although several human and animal studies have demonstrated a relationship between bisphenol exposure and insulin resistance,^10^ the low dose of BPS used in the present study compromised microvascular endothelial function without influencing glucose homeostasis. These findings highlight that the endothelium of the microvasculature may be particularly susceptible to bisphenol exposure Using an *in vitro* HUVEC model, we show that exposure to BPS as low as 2.5 nM induces increased production of ROS. These findings are consistent with other studies showing that excess ROS production in multiple cell types arises from exposure to BPA or its analogues, suggesting that oxidative stress may be an important cellular mechanism that links bisphenol exposure to various diseases and disorders. Vascular oxidative stress decreases the availability of NO and is a common cause of microvascular endothelial dysfunction.^17,26^ When the production of O_2_^−^ in the vasculature exceeds antioxidant defenses, O_2_^−^ reacts with NO, reducing is bioavailability and leading to the production of peroxynitrite (ONNO^−^), a highly reactive molecule that oxidizes macromolecules, contributing to cellular dysfunction. Oxidative degradation of BH4, a critical cofactor that stabilizes the active dimeric form of eNOS and facilitates electron transfer for the oxidation of L-arginine, is thought to contribute to uncoupling in which eNOS produces more O_2_^−^ than NO.^27^ Uncoupling of eNOS fuels a cycle of excess ROS production, oxidative stress and reduced NO availability. In HUVEC exposed to BPS, NO production was suppressed, while supplementation of BH4 or inhibition of eNOS normalized ROS production, suggesting that uncoupled eNOS contributes to BPS-induced oxidative stress in endothelial cells. Thus, reduced NO bioavailability due to vascular oxidative stress may be the mechanism underlying impaired endothelium-dependent vasodilation.

In addition to uncoupled eNOS, vascular ROS arise from other enzymatic sources, including xanthine oxidase and the NADPH oxidases, as well as the mitochondria electron transport chain. Under normal conditions, electrons are leaked from redox centres of the electron transport chain, reducing molecular O_2_ to the superoxide anion, O_2_^−^, which is quickly converted to H_2_O_2_ by mitochondria SOD and subsequently reduced to water by glutathione peroxidase. Mitochondria are not only sources of ROS, but also targets of ROS that are prone to oxidative damage. Dysfunctional mitochondria produce excess ROS that overwhelm antioxidant defenses and contribute importantly to the development of atherosclerosis and other CVD.^28,29^ Our findings show increased production of mitochondria-derived ROS in BPS-exposed HUVEC and restoration of redox balance and NO availability in HUVEC treated with mitochondria-targeted antioxidants prior to BPS exposure, suggesting that dysfunctional mitochondria contribute to the instability of the redox environment of endothelial cells. Bioenergetic profiling revealed no perturbations in basal or maximal mitochondrial respiration after 48 hrs of exposure, except for a decrease in respiratory reserve capacity. However, using both the TMRE and JC-1 assays, we show that BPS exposure results in a loss of mitochondria membrane potential, which is associated with increased ROS production by mitochondria and triggers mitophagy and cell death.

The adverse health effects of bisphenols have largely been attributed to their endocrine disrupting properties; however, our data reveal oxidative stress to be an important mechanism linking bisphenols to cardiovascular dysfunction. It is possible that the endocrine disrupting properties of BPS modulate the redox response, and this may explain the sex-dependent effects we observed in the microvasculature of BPS-exposed mice. BPA and its analogues are known to bind to estrogen receptors (ER), albeit with lower affinity than endogenous estrogen, and exhibit estrogenic activity. In endothelial cells, ERα localizes to caveolae in the cell membrane and activates Akt-dependent NO production via phosphorylation of eNOS; while the transcription of eNOS is activated upon binding of cytosolic or nuclear ERα/ERβ to estrogen response elements in the eNOS promoter.^30^ Acute activation of ERα leads to NO-dependent vasodilation and suppression of O_2_^−^ levels, including mitochondria-derived O_2_^−^.^31^ The use of a HUVEC line in the current study precluded investigation into the molecular basis of sex differences observed in BPS-exposed mice; however, data from our lab show that acute exposure to BPS augments NO-dependent relaxation in isolated aortae from female mice only. These findings suggest that the estrogenic activity of BPS counteracts its ROS-promoting effects, providing a potential explanation for the sex-dependent vascular effects observed in the present study where females appeared protected from BPS-induced endothelial dysfunction.

## Summary

The current study demonstrates that chronic exposure to a low dose of the commonly used BPA substitute, BPS, leads to sex-dependent impairments in NO-mediated vasodilation in the microvasculature. Our findings substantiate epidemiological evidence of a link between bisphenol exposure and vascular disease, by providing direct evidence of endothelial dysfunction in pressurized microvessels; however, interpretations are limited to a single dose and single toxicant exposure. Using a HUVEC cell line that preserves characteristics of primary endothelial cells, we provide evidence that BPS-induced endothelial dysfunction is due to NO depletion as a result of eNOS uncoupling, mitochondria dysfunction and consequent redox imbalance. Thus, our study highlights a potential molecular mechanism underlying the link between bisphenol exposure and vascular diseases observed in human populations.

## Figure Legends

**Supplementary Figure 1:** Representative tracings from pressure myography. After equilibration of vessels at 60 mmHg and testing of viability, mesenteric arteries from control (A) or BPS-exposed males (B) were pre-contracted with Phe and dilated with cumulative concentrations of Mch. After washing, contractile responses to cumulative doses of Phe were assessed. Internal diameter was recorded and normalized to passive diameter in Ca^2+^-free conditions.

**Supplementary Figure 2:** Glucose tolerance in mice exposed to BPS postnatally. Glucose tolerance tests were performed in fasted male (A) or female (B) C57BL/J6 mice after exposure to 250 nM BPS via drinking water. Glucose was measured with a glucometer at baseline and after injection of glucose (IP). n = 9-14 mice/group.

## Acknowledgements

The salary of RDS was supported by fellowships from the Molly Towell Perinatal Research Foundation and the Libin Cardiovascular Institute. The salaries of TS and LC were supported by scholarships from the Libin Cardiovascular Institute. LC also received scholarships from the Cumming School of Medicine and Faculty of Graduate Studies at the University of Calgary. LB received salary support from the University of Calgary. The research performed for this study was supported by funding from the Libin Cardiovascular Institute and the Canadian Foundation for Innovation (CFI). JAT was supported by a National New Investigator Award from the Heart and Stroke Foundation of Canada (HSFC). The authors thank Dr. Andy Braun for his feedback on the manuscript and Dr. Steven Greenway for kindly providing the SS31 mitochondria-targeted antioxidant.

