## Supplementary figures and images for "Exploring oxidative stress and endothelial dysfunction as a mechanism linking bisphenol S exposure to vascular disease in human umbilical vein endothelial cells and a mouse model of postnatal exposure"

### Supplementary Figure 1

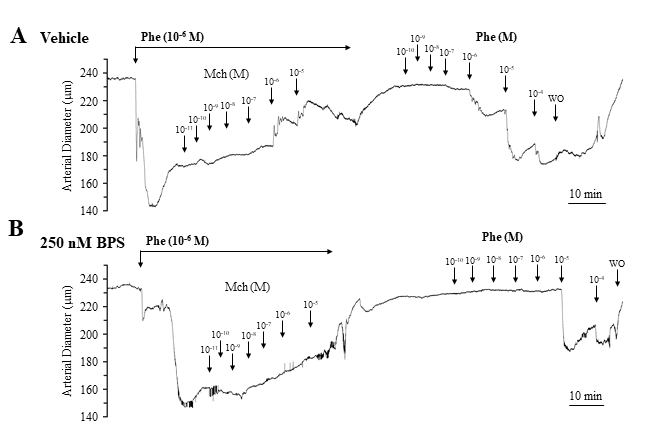

### Supplementary Figure 2

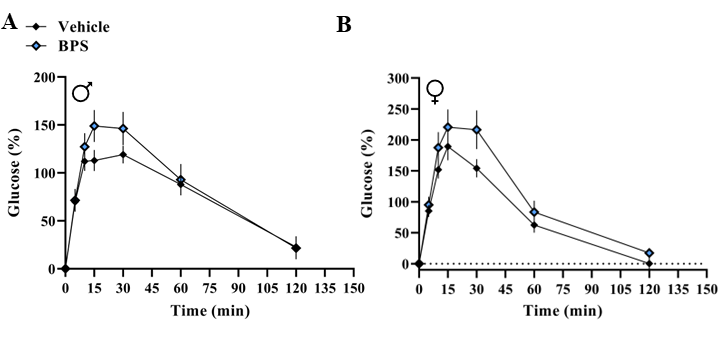
